# Towards the minimal proteome of life: Proteome profiles of *Bacillus subtilis* vegetative cells and spores

**DOI:** 10.1101/678979

**Authors:** Bhagyashree Swarge, Winfried Roseboom, Chris G. de Koster, Stanley Brul, Leo J. de Koning

**Affiliations:** Swammerdam Institute of Life Sciences, University of Amsterdam

## Abstract

The method of ^15^N metabolic labelling of *Bacillus subtilis* enabled mass spectrometric quantification of relative protein levels in the vegetative cells and the spores of this model organism. A total of 1501 proteins have been identified from the combined spore and vegetative cell samples. From these 1086 proteins have been relatively and reproducibly quantified between spores and vegetative cells. Of the quantified proteins, 60% are common to the vegetative cells and spores, indicating that spores host a minimal set proteins sufficient for the resumption of metabolism upon completion of germination. The shared proteins represent, the most basic ‘survival kit’ for life on earth that is known thus far.

## Introduction

Endospore formation is characterized by a continues protein turnover and by various cellular rearrangements^1,2^. During protein turnover existing proteins are degraded and new proteins are formed. These processes are controlled by sporulation specific sigma factors and involve various cellular structural rearrangements such as asymmetric cell division and the extrusion of water from the developing endospore^2^. The reorganization involves a macromolecular machinery and results in the formation of multi-layered spores that are resistant to UV, radiation, heat and different chemical agents^3^. In addition to the development of stress resistance, spores also prepare to equip themselves with all necessary elements that are essential to return to life. These elements generally are comprised of proteins that steer germination and outgrowth to the point where cells resume their vegetative cell cycle. To jump start the germination, the macromolecules needed are pre-synthesized during sporulation^4^. Even though the spores are metabolically dormant^5^ researchers are still in pursuit of finding the exact molecular, biochemical and biophysical mechanisms behind the germination process^6−8^. Despite numerous indications from genetic studies, the existing hypotheses on the molecular basis of spore revival remains, at the mechanistic level, unanswered^8^. Processes such as water uptake, dipicolinic acid (DPA) release, cortex hydrolysis and signal transduction therein are still not completely understood^5−7^. All these processes are mediated via a number of different proteins^9^. Many proteomics and transcriptomics studies^10−13^ have addressed the set of genes and their inferred proteinaceous counterparts in an effort to make a comprehensive inventory of the putative proteins mediating the described physiological processes. Yet, none of them has systematically focused on a comparative, quantitative, direct proteome analysis of dormant spores versus the vegetative cells into which they transform.

In the present study, we have quantitatively characterized the *B. subtilis* spore proteome relative to that of vegetative cell. To achieve this, spores are mixed with ^15^N-metabolically labelled vegetative cells and the mixture is processed with our recently developed one-pot method^14^ for mass-spectrometric analyses. Here, the ^14^N/^15^N isotopic protein ratios represent the relative spore over vegetative cell protein abundances. The mass spectrometric relative quantification has led to the quantification of over 1000 proteins from combined spore and cell samples. We aim to deduce the differences in the two proteomes and classify a basic set of proteins harboured within a dormant spore that constitutes a minimal set of proteins sufficient for the life to survive severe environmental stresses and resume growth when conditions are again more favourable.

## Materials and methods

### Bacterial strain, media, and culturing conditions

*B. subtilis* wild-type strain PY79 was used for preparing ^14^N (Light) spores and ^15^N (Heavy)-labelled reference vegetative cells. For sporulation, bacteria were pre-cultured and sporulated as described previously^14,15^. Defined minimal medium, buffered with 3-(N-morpholino) propane sulfonic acid (MOPS) to pH 7.4 was used for sporulation^16^. The spore cultures were grown and sporulated in the presence of ^14^NH_4_Cl while the reference vegetative cell cultures received ^15^NH_4_Cl as the sole nitrogen source. The final stock of reference vegetative cells consisted of cells harvested at exponential growth. The cells were analysed against three independent biological replicates of spore query cultures.

### Mixing of ^14^ N spores and ^15^N-labeled cells, and one-pot sample processing

The harvested ^14^N-spores were mixed in a 1:1 ratio (based on the cell count) with ^15^N-labelled vegetative cells. After mixing, the samples were further subjected to one-pot sample processing as described previously^14^. Typically, a mixture of spores and cells was suspended in lysis reduction buffer and disrupted in seven cycles with 0.1 mm zirconium-silica beads (BioSpec Products, Bartlesville, OK, USA) using a Precellys 24 homogenizer (Bertin Technologies, Aix en Provence, France). The tubes were incubated for 1 h at 56°C and alkylated using 15 mM iodoacetamide for 45 min at room temperature in the dark. The reaction was quenched with 20 mM thiourea and the first digestion with Lys-C (at 1:200 protease/protein ratio) was carried out for 3 h at 37°C. Samples were diluted with 50 mM ammonium bicarbonate buffer and the second digestion with trypsin (at 1:100 protease/protein ratio) was carried out at 37°C for 18 h. The tryptic digest was freeze-dried. Before use, the freeze-dried samples were re-dissolved in 0.1% Tri-fluoric acid (TFA) and cleaned up using C18 reversed phase TT2 TopTips (Glygen), according to the manufacturer’s instructions. The peptides were eluted with 0.1% TFA in 50% acetonitrile (ACN) and freeze dried.

### Fractionation of peptides in one-pot digests

A ZIC-HILIC chromatography was used to fractionate the freeze-dried peptide samples. Dried digests were dissolved in 200 µl of Buffer A (90% acetonitrile, 0.05% Formic acid, pH 3), centrifuged to remove any undissolved components, and injected into the chromatography system. An isocratic flow with 100% Buffer A for 10 min was followed by a gradient of 0-30% Buffer B (25% acetonitrile, 0.051% formic acid, pH 3) in the first phase and 30-100% of Buffer B in the second phase (flow rate 400 µl/min). The peptides were eluted and collected in 10 fractions, freeze-dried, and stored at −80°C until further use.

### LC-FT-ICR MS/MS analysis

ZIC-HILIC fractions were re-dissolved in 0.1% TFA and peptide concentrations were determined by measuring the absorbance at wavelength of 215 nm with a NanoDrop. LC-MS/MS data were acquired with an Apex Ultra Fourier transform ion cyclotron resonance mass spectrometer (Bruker Daltonics, Bremen, Germany) equipped with a 7 T magnet and a Nano electrospray Apollo II Dual Source coupled to an Ultimate 3000 (Dionex, Sunnyvale, CA, USA) HPLC system. Samples containing up to 300 ng of the tryptic peptide mixtures were injected as a 10μl 0.1% TFA, 3% ACN aqueous solution together with 25 fmol of [Glu1]-Fibrinopeptide B human peptide and loaded onto a PepMap100 C18 (5 μm particle size, 100 Å pore size, 300 μm inner diameter × 5 mm length) precolumn. Following injection, the peptides were eluted through an Acclaim PepMap 100 C18 (3 μm particle size, 100 Å pore size, 75 μm inner diameter × 250 mm length) analytical column (Thermo Scientific, Etten-Leur, the Netherlands) to the Nano electrospray source. Gradient profiles of up to 120 min were used from 0.1% formic acid-3% acetonitrile to 0.1% formic acid-50% acetonitrile (flow rate 300 nl/min). Data-dependent Q-selected peptide ions were fragmented in the hexapole collision cell at an argon pressure of 6 × 10^−6^ mbar (measured at the ion gauge) and the fragment ions were detected in the ICR cell at a resolution of up to 60,000. In the MS/MS duty cycle, three different precursor peptide ions were selected from each survey MS. The MS/MS duty cycle time for one survey MS and three MS/MS acquisitions was approximately 2s. Instrument mass calibration was better than 1 ppm over m/z range of 250– 1500.

### Data analysis and bioinformatics

Each raw FT-MS/MS data set was mass calibrated better than 1.5 ppm on the peptide fragments from the co-injected GluFib calibrant. The 10 ZIC-HILIC fractions were jointly processed as a multifile with the MASCOT DISTILLER program (version 2.4.3.1, 64 bits), MDRO 2.4.3.0 (MATRIX science, London, UK), including the Search toolbox and the Quantification toolbox Peak-picking for both MS and MS/MS spectra were optimized for the mass resolution of up to 60000. Peaks were fitted to a simulated isotope distribution with a correlation threshold of 0.7, with minimum signal to noise ratio of 2. The processed data were searched in a MudPIT approach with the MASCOT server program 2.3.02 (MATRIX science, London, UK) against the *B. subtilis* 168 ORF translation database. The MASCOT search parameters were as follows: enzyme - trypsin, allowance of two missed cleavages, fixed modification - carboamidomethylation of cysteine, variable modifications - oxidation of methionine and deamidation of asparagine and glutamine, quantification method – metabolic ^15^N labelling, peptide mass tolerance and peptide fragment mass tolerance - 50 ppm. MASCOT MudPIT peptide identification threshold score of 20 and FDR of 2% were set to export the reports.

Using the quantification toolbox, the quantification of the light spore peptides relative to the corresponding heavy cell peptides was determined as ^14^N/^15^N ratio using Simpson’s integration of the peptide MS chromatographic profiles for all detected charge states. The quantification parameters were: Correlation threshold for isotopic distribution fit - 0.98, ^15^N label content - 99.6%, XIC threshold - 0.1, all charge states on, max XIC width −120 seconds, elution time shift for heavy and light peptides - 20 seconds. All Isotope ratios were manually validated by inspecting the MS spectral data. The protein isotopic ratios were then calculated as the average over the corresponding peptide ratios. For each of the three replicas the identification and quantification reports were imported into a custom-made program to facilitate data combination and statistical analysis. Protein identification was validated with identifications in at least two replicas. For these identified proteins the relative quantification was calculated as the geometric mean of at least two validated ^14^N/^15^N ratios. All identification and quantification protein data are listed in **Supplementary Table 1** with the standard errors and geometric standard deviations. DAVID Bioinformatics Resources tool (version 6.8) was used^17^ to retrieve the data of UniProt key word and KEGG pathway classifications. Data are available via ProteomeXchange with identifier PXD014335. Membrane proteins were predicted (**Supplementary Table 2)** using the following programs or databases: PSORTb 3.0 (http://www.psort.org/psortb/), Phobius (http://phobius.sbc.su.se/), BOMP (http://services.cbu.uib.no/tools/bomp), PRED-LIPO (http://bioinformatics.biol.uoa.gr/PRED-LIPO/), LocateP (http://www.cmbi.ru.nl/locatep-db/cgi-bin/locatepdb.py), and InterProScan 5 (http://www.ebi.ac.uk/interpro/). Identified proteins were categorized according to SubtiWiki (http://subtiwiki.uni-goettingen.de/).

## Results

### Identification and quantification of the vegetative cell and spore proteins

A total of 1501 proteins are identified from the combined spore and vegetative cell (henceforth referred to as cell) samples, of which 1086 are relatively and reproducibly quantified. Based on the average protein signal/noise (S/N) ratio we operationally defined predominant spore and predominant vegetative cell proteins. Thus, isotopic ^14^N/^15^N ratios of more than 20 predominantly correspond to spore proteins and isotopic ^14^N/^15^N ratios of less than 0.05 predominantly correspond to vegetative cell proteins. The isotopic ratios 20> ^14^N /^15^N >0.05 correspond to the proteins shared between the spores and cells. **Figure 1** represents a distribution of the quantified proteins, where the combined amounts of the spore and vegetative cell proteins indicated by the log_10_ values of their MASCOT scores are plotted against the corresponding relative protein levels indicated by the log_2_ values of the experimental ^14^N/^15^N ratios. In this study, 130 proteins are found to be spore-predominant, while 299 proteins are considered to be cell-predominant. From the remaining 657 shared proteins, only 7% are enriched in spores, with ^14^N/^15^N ratios between 1 and 20, while 93% are enriched in cells, with ^14^N/^15^N ratios between 1 and 0.05.

**Figure 1.**
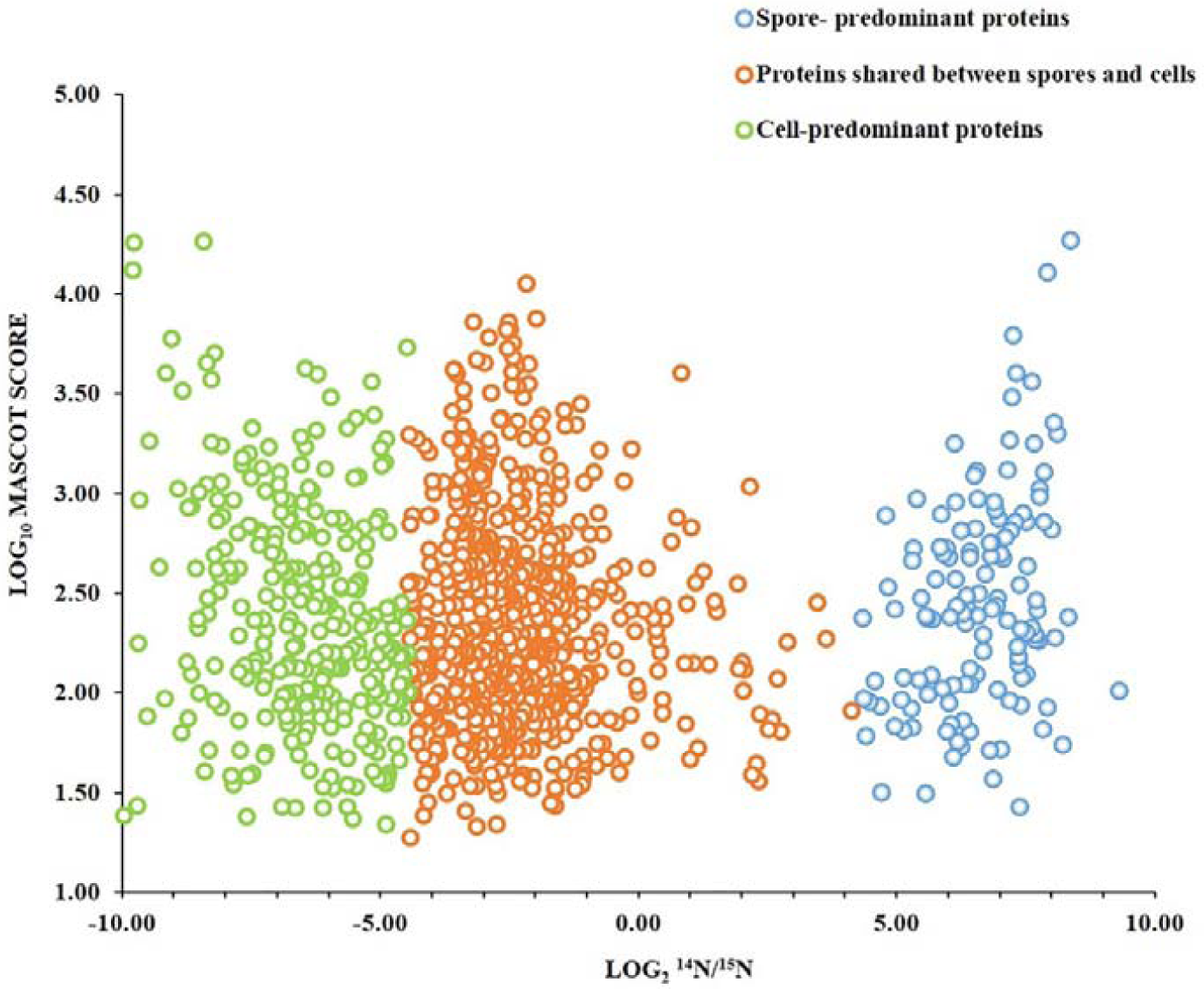
Distribution of proteins in *Bacillus subtilis* spores and vegetative cells. MASCOT scores indicate the combined spore and cell abundance of a protein versus its ^14^N/^15^N protein isotopic ratio, which represents the relative level of the protein in the spores over the vegetative cells. The blue circles indicate spore-predominant proteins (^14^N/^15^N > 20), green circles indicate vegetative cell-predominant proteins (^14^N/^15^N < 0.05), and orange circles indicate proteins shared between spores and vegetative cells (20 > ^14^N/^15^N > 0.05).

### Spore-predominant proteins

Most of the proteins belonging to this category are germination and structural spore proteins. These include spore coat proteins, small acid soluble proteins (SASPs) being most abundant, and germination proteases. Interestingly, 24 hydrolases are detected, which include the protein YyxA with the highest levels in spores **(Supplementary Table 1)**. It is noteworthy that of all the metabolic pathway enzymes, only MalS is present in abundant quantities in spores (**Supplementary Figure 1**). The bioinformatic analysis has predicted 31 spore predominant membrane proteins of which proteins CoxA, YhcN, YdcC, GerD, YlaJ, all associated with germination, are classified as lipid-anchored proteins.

### Cell-predominant proteins

Cell surface proteins belonging to the surfactin family are found to be the most abundant **(Supplementary Table 1)** in this group. Next to it, proteins involved in purine and pyrimidine biosynthesis are exclusively found in cells. In addition, branched chain amino acid synthesis, leucine, threonine, thiamine and methionine biosynthesis proteins are predominant in cells **(Table 1)**. Interestingly, most of the tricarboxylic acid (TCA) cycle enzymes are present in the cell-predominant category (**Supplementary Figure 1**). Membrane prediction analysis has classified 93 proteins as cell-predominant membrane proteins.

**Table 1.** Uniprot keywords annotation enrichment of quantified *Bacillus subtilis* PY79 spore and vegetative cell proteins based on DAVID functional annotation analysis.

| UniProt Keyword<sup>a</sup> | Number of proteins |  |  |
| --- | --- | --- | --- |
|  | Spore proteome | Vegetative cell proteome | Shared between cells and spores |
| Sporulation | 77 |  |  |
| Peroxidase | 6 |  |  |
| Carboxypeptidase | 5 |  |  |
| Ribosome biogenesis | 7 |  |  |
| Cytoplasm | 193 | 243 | 186 |
| Ribosomal protein | 49 | 49 | 49 |
| Protein synthesis | 36 | 36 | 36 |
| Nucleotide binding | 134 | 189 | 131 |
| Oxidoreductase | 96 | 112 | 85 |
| ATP synthesis | 7 | 7 | 7 |
| Hydrolase | 123 | 136 | 103 |
| Fatty acid biosynthesis | 10 | 11 | 10 |
| Cell cycle | 14 | 22 | 14 |
| Cell Division | 14 | 22 | 14 |
| Amino acid biosynthesis | 24 | 75 | 24 |
| NAD | 38 | 45 | 37 |
| Zinc | 52 | 56 | 47 |
| NADP | 33 | 40 | 32 |
| Lipid biosynthesis | 14 | 12 | 15 |
| rRNA processing | 10 | 11 | 10 |
| Kinase | 32 | 45 | 32 |
| Cell shape | 17 | 16 | 15 |
| RNA-binding | 59 | 60 | 59 |
| Stress response | 33 | 41 | 33 |
| Transferase | 101 | 158 | 101 |
| rRNA-binding | 35 | 35 | 35 |
| Isomerase | 28 | 34 | 27 |
| Lysine biosynthesis | 8 | 10 | 8 |
| tRNA-binding | 11 | 11 | 11 |
| Topoisomerase | 5 | 5 | 5 |
| Aminoacyl-tRNA synthetase | 21 | 21 | 21 |
| Protein transport |  | 14 |  |
| Nucleotide biosynthesis |  | 25 |  |
| Threonine biosynthesis |  | 4 |  |
| Methionine biosynthesis |  | 16 |  |
| Histidine biosynthesis |  | 9 |  |
| Arginine biosynthesis |  | 9 |  |
| Leucine biosynthesis |  | 4 |  |
| Threonine biosynthesis |  | 4 |  |
| Branched chain amino acid<br>biosynthesis |  | 13 |  |
| Thiamine biosynthesis |  | 11 |  |
| DNA replication |  | 12 |  |
| Septation |  | 10 |  |
| Flagellar rotation |  | 5 |  |
| Chemotaxis |  | 16 |  |
| Methylation |  | 9 |  |
| Flavoprotein |  | 29 |  |
| Multifunctional enzyme |  | 20 |  |
| Iron-sulfur |  | 20 |  |
| Allosteric enzyme |  | 7 |  |

<sup>a</sup>EASE score i.e. *p*-value threshold for the keyword annotation enrichment was set to 0.05

### Proteins shared between spores and cells

The proteins shared between spores and cells are mostly ribosomal proteins, cell cycle-regulating and/or associated proteins, and cytosolic proteins involved in the pathways required for anabolism and catabolism of proteins, carbohydrates, lipids and pathways of energy metabolism. These proteins are organized in 50 different categories by DAVID^17,18^ **(Table 1)**. Many of the proteins encoded by essential genes are also observed to be shared. These include tRNA-synthetases, carboxylases involved in metabolism, DNA polymerases and RNA processing as well as degradation proteins. Only 20% of the proteins involved in amino acid biosynthesis are present in spores and a majority is enriched in cells (**Table 1, Figure 2**). **Figure 2** represents the averaged levels of proteins belonging to latent metabolic pathways and related functional categories that are observed in spores. Many of the proteins from these pathways and functional categories are needed to start the germination. As exemplified in **Supplementary Figure 1**, these shared proteins indicate the metabolic options available to the spores when they initiate germination. In conclusion, this category of proteins substantially contributes to the basic set of proteins of a dormant spore. More than 130 proteins from this group are found to be membrane proteins.

**Figure 2.**
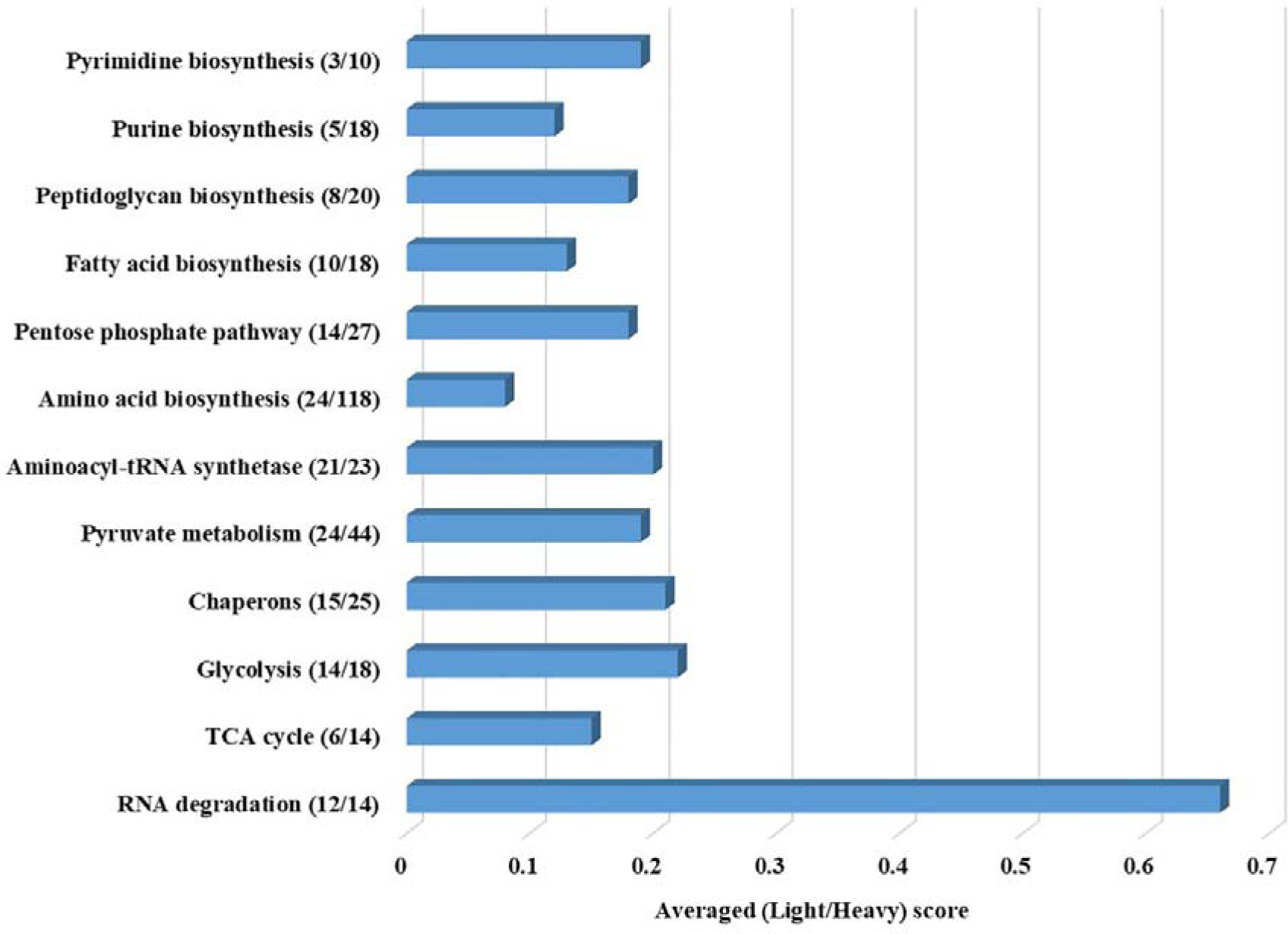
Latent metabolic pathways and related functional categories in the dormant spores. X-axis represents averaged ^14^N/^15^N ratios of proteins belonging to respective pathways indicating their levels in spores. The numbers in the parenthesis represent the numbers of the quantified shared category proteins / the total number of theoretical proteins belong to the pathways according to DAVID.

## Discussion

Dormant spores of *B. subtilis* are metabolically inactive and their main function is to protect the genetic information throughout the period of unfavourable environmental conditions. Subsequently, a spore must be able to germinate and allow vegetative cells to proliferate and populate a new environmental niche. Here, we discuss the proteomes of the spores and the vegetative cells that are studied, to our knowledge, for the first time in a single experiment. We aim to understand the fundamental differences between these two forms of the spore forming organism *Bacillus subtilis*. Metabolic labelling using ^15^N isotope is an easy and highly accurate means of proteome quantification^19^. Combining the metabolic labelling approach with our one-pot sample processing method^14^ has shown that only 40% of the quantified proteins make up both the spore-predominant and cell-predominant categories. The majority of the proteins classified as spore-predominant, are structural proteins such as spore coat proteins, cortex lytic enzymes and SASPs. Whereas surfactin synthases, flagellar proteins (FliT, FlhP, FlgG), cell division protein FtsZ, SepF, FtsA, DivIVA and the proteins involved in amino acid biosynthesis are found to be predominant in vegetative cells. The remaining 60% of the quantified proteins are shared between spores and cells. The presence of enzymes for macromolecular synthesis and energy metabolism in dormant spores is well studied^20,21^ and our data shows that the corresponding pathways are shared between cells and spores albeit that the relative quantities of the proteins vary in both. A discussion of quantified and functionally vital proteins of *B. subtilis* is presented below. This represents a basic set of proteins in a dormant spore that awaits germination cues and as such is a ‘survival kit’ of life on earth.

### The Phosphoenolpyruvate dependent phosphotransferase system (PTS)

PTS is the carbohydrate uptake system in *B. subtilis*^22^. We find that the components of glucose transport are present in dormant spores. Glucose is one of the germinants triggering germination and it is most likely transported into the spore by PtsG, a transmembrane protein, which is present in the spores. The uptake further likely involves the central proteins HPr (PtsH) and Enzyme I (PtsI). In addition, the transporters for fructose (FruA) and sucrose (SacP) are also identified in our spore samples. Although glucose is a preferred source of carbon over fructose, the levels of FruA are higher than those of PtsG in spores. In contrast to these PTS proteins, the putative glucosamine transporter (GamP) is abundantly present only in cells.

### Other transporters

In our data almost 40 transporter proteins, have been quantified. These proteins belong to the ABC transporters family, including the protein and amino acid transporters, as well as families of ion transporters. Interestingly, in a recent transcriptome study, almost 80 transcripts encoding different transporters have been identified from spore mRNA isolates^23^. The ABC transporters, functioning in the import or export of amino acids, peptides and polysaccharides^24^ might play key roles in both sporulation and germination processes. Protein AtcL (YloB), a calcium transporting ATPase found in high levels in spores, facilitates uptake of Ca^2+^ during sporulation^25^. During germination different germinants such as amino acids, sugars and cations are transported via different transporters through the coat and the cortex layers to various germinant receptors located in the spore inner membrane. GerPC, a spore predominant protein, is one such transporter known to facilitate the transport of germinants to their cognate GRs^26^. TcyA, a cysteine binding transporter, quantified as a shared protein, is functionally similar to YckB (identified only in one replicate) whereas YtnA, an amino acid permease, and ArtM transporting arginine are also shared between spores and cells. EcsA,a regulator of protein secretion^27^, is essential in sporulation^24^ and might play a role in germination due to its inherent protein transporting capacity. The uncharacterized proteins YvrC, YkpA, YdbJ, YfmR, YknZ, YhaQ, YfmM and YhfQ from ABC transporter family are also shared by spores and cells of which YhfQ is has been linked to siderophore uptake^28^. The *B. cereus* orthologs of YfmM and YdbJ (BC5433 and BC1359 respectively), as well as of YknZ (BC5253) are reported to be functional drug efflux pumps, more specifically for the latter is a macrolide exporter^29^. Also the orthologs of YhaQ, BerA in *B. thuringiensis* and BcrA in *B. licheniformis* are concerned with β-exotoxin^30^ and bacitracin^31^ resistance respectively. Proteins YkpA and YmfR, being paralogs of YfmM, thus could also be involved in drug efflux. These findings highlight the potential role of the ABC transporters with respect to *B. subtilis* spore resistance properties. In the cell-predominant category, we have quantified MetN that imports methionine and YufP, YhdG, AlsT that function as amino acid permeases. YufP is an uncharacterized ABC transporter protein, predominantly quantified in cells that may act as a branched chain amino acid transporter protein. Out of the five osmoprotectant transporters (OpuA-E) only OpuA transporters (OpuAA, OpuAB, OpuAC) are quantified in spores; whereas, OpuB and OpuC transporters (OpuBA, OpuBC, OpuCA, OpuCC) are quantified predominantly in cells. The Opu family of transporters are known for the uptake of glycine betaine.

### Proteases and peptidases

From the group of catalytic enzymes, the proteases and peptidases play a crucial part in spore formation, its resistance, germination as well as in stress survival of vegetative cells. In this study, 36 proteases and 16 peptidases are quantified. These, amongst others, include germination protease (Gpr), and DD-carboxypeptidase involved in spore wall maturation. In addition to that, the proteases involved in cellular regulatory processes include ATP dependant Clp proteases, metalloproteases, Lon proteases, and aminopeptidases. Protein YyxA, a predictive Htr-type serine protease is suggested to play important role in spore germination by interacting with SleB^32−34^ and sporulation specific protease YabG, is involved in coat protein modification and cross linking of coat proteins such as GerQ^33,35^. Both these proteins are observed to be spore predominant in current strudy. CwlC, CwlJ, SleB are spore-predominant cell wall hydrolases. CwlC functions similar to LytC (CwlB) in mother cell lysis during sporulation whereas the other two lytic enzymes play an important role in cortex hydrolysis during germination^36,37^. Among the cell-predominant group, the cell wall lytic enzymes LytF, LytE and LytC are observed in large quantities. Of these, LytF and LytE are endopeptidases having crucial roles in cell separation during cell division whereas LytC is an amidase that plays an important role in the hydrolysis of the N-acetylmuramoyl-l-alanine linkage in peptidoglycan^38,39^. The enzyme thus plays an important role in cell separation^40^ and mother cell wall lysis during spore formation^41^. Cell predominant extracellular protease, WprA, is a cell surface protease ^42^. This protein is expressed during exponential growth of *B. subtilis* ^42^ and is needed for stabilization of lytic enzymes and septation^39^.

### Protein synthesis

To survive in harsh environments, bacteria have a minimal set of essential genes. *Bacillus subtilis* has total of 261 essential genes^43^ in its genome. These genes are involved in information processing, synthesis of components necessary for maintaining the cell shape, synthesizing the envelope and cell energetics^44^. The largest category belongs to the essential genes related to protein synthesis. In our data set, except the 30S ribosomal protein S18 (RpsR) and 50S ribosomal protein L19 (RplS) all the essential as well as non-essential ribosomal proteins are identified in both the vegetative cells and spore. However, individual levels of these ribosomal proteins in a spore have dropped to about 20% of their levels in a vegetative cell (**Supplementary Table 1**). Nevertheless, the levels of 50S ribosomal protein L31 type B (RpmE2) in spores are comparable to those in cells. It has been demonstrated that in *B. cereus* the spore ribosome are damaged and distinctively different from those of vegetative cells^45^. On the contrary, the spore ribosomes in *B. megaterium* are similar to those of the vegetative cells and the activity of these proteins is not altered during sporulation^46,47^. The high levels of ribosome maturation factors RimM and RimP in spores may help in efficient production of translationally competent ribosomes^48^ during germination. In addition to the ribosomal proteins, the aminoacyl tRNAs found in the spore proteome may provide a ‘ready to use kit’ for the amino acid biosynthesis and thereby protein synthesis during germination^49^.

### Amino acid metabolism

Amino acid degradation plays an important role during germination as well as cell homeostasis of outgrowing cells. In dormant spores, the amino acids lysine, arginine and glutamate are relatively abundant and their levels are elevated immediately after germination as compared to those of other amino acids^50^. The presence of the enzymes arginase (high levels in dormant spores) and leucine dehydrogenase (levels comparable to those in vegetative cells) in spores corroborate the thesis that the respective amino acids may have a role in the progression of the spore germination process. Glutamate dehydrogenase (RocG), though found at very low levels in spores, may play a role in glutamate utilization^51^ along with the Gpr (germination protease) acting on SASPs. Moreover, arginine kinase in spores, phosphorylates arginine and thus marks proteins to be cleaved by Clp proteases^52^. Two enzymes 1-pyrroline-5-carboxylate dehydrogenase (RocA) and 1-pyrroline-5-carboxylate dehydrogenase 2 (PutC) are shared in spores and vegetative cells and are quantified with equal levels in both phenotypes. These are involved in the catabolism of proline and arginine to glutamate^53,54^. These enzymes are present in the mother cell during sporulation and thus may get entrapped in or are associated with the spore^53^. The components BfmBAB and BfmBB of the branched-chain 2-keto acid dehydrogenase complex (BDCH), are quantified in the shared category and are involved in valine and isoleucine utilization to form branched chain fatty acids^55^. The third component BfmBAA is identified only once and thus not quantified. Very few proteins involved in the biosynthesis of amino acids such as methionine (MtnN, FolD, MetA, Asd), lysine (DapF, DapA, DapH, YkuR, LysA, PatA) and histidine (HisIE, HisA, HisD, HisB) are present in the spores. One single protein is quantified for the biosynthesis of aromatic amino acid, branched chain amino acid, proline and threonine. The remaining proteins involved in these processes are dominantly recovered in vegetative cells.

### Metabolic pathways and Energy production

From the quantified data set, 290 proteins are classified to the category of metabolic pathways by DAVID functional annotation. These proteins belong to carbon metabolism pathways including glycolysis, pentose phosphate pathway and the TCA cycle as well as to pyruvate metabolism, amino acid biosynthesis, purine and pyrimidine metabolism, fatty acid biosynthesis, peptidoglycan biosynthesis. In our data, many glycolysis and pentose phosphate pathway proteins related to energy production are found to be present in spores. The fact that the enzyme glucose dehydrogenase (Gdh) is produced in the forespore during later stages of sporulation ^56^ may explain the observed high levels of this enzyme in the spores. It has been observed in the past, that this enzyme reacts with glucose in presence of NAD^+^ and NADP^+56^ which are also already present in the spores in high amounts^57^. Thus during germination, glucose dehydrogenase together with the phosphogluconate dehydrogenase (GndA) may fuel the glucose (available as a germinant) into the pentose phosphate pathway^58,59^. Interestingly, only two enzymes of the TCA cycle, fumarase and isocitrate dehydrogenase appear to be shared between the spores and the cells whereas other enzymes of the pathway are exclusively found in cells. This is in agreement with the study in *B. thuringiensis* where the level of the TCA cycle proteins are significantly reduced during sporulation^60^. It has also been observed that the levels of α-ketoglutarate dehydrogenase, succinate dehydrogenase, and malate dehydrogenase are decreased at stationary phase whereas the levels of fumarase and malic enzymes remain high^61^ in *B. subtilis*. Interestingly, malic enzyme MalS is highly abundant in spores, as mentioned previously while the other malic enzymes YtsJ and YqkJ are present in both spores and cells. A recent study suggested that L-malate stores in the spore are used as energy source during germination. In support of this MalS is found to be one of the early proteins synthesized in spore^12^. Since malate and glucose are preferred carbon sources for *B.subtilis*^62^, it is plausible that the abundant levels of MalS can be utilized by the spore during germination if malate is available in the environment. However, a recent study has observed that the levels of L-malate in spores are very low casting significant doubt on its proposed role in energy generation^63^. Instead the enzyme might be involved in maintaining redox balance in the initial phases of spore outgrowth by synthesizing malate at the expense of reduction equivalents (NADH) thus potentially contributing to NAD regeneration for a sustained rapid catabolism of 3-phosphoglyceric acid, a prime energy reserve stored in spores^63^.

In dormant spores of *B. subtilis*, levels of ATP are less than 1% of those in the cells and the ATP generation is suggested to resume only after germination is complete^64^. In *B. thuringiensis*, the expression of F_0_F_1_-ATPase subunits is upregulated during sporulation in order to keep a constant energy supply to the sporulating cell^60^. Remarkably, we observe increased levels of seven F_0_F_1_-ATP synthase subunits in our data. All of these subunits are associated with either the protein module that synthesizes/hydrolysizes ATP or with the module involved in proton transport^65^. Adenylate kinase, a phosphotransferase enzyme, present in the shared group of proteins, plays an important role in maintaining levels of adenine nucleotides. It has been observed that the level of adenylate kinase in spores are only about 10% of the levels found in early stationary-phase cells^66^. Moreover, for the purine biosynthesis pathway, the proteins involved in the conversion of inosine monophosphate (IMP) to ATP (GuaB, GuaA, Ndk) and IMP to GTP (PurA, Adk) are shared between the spores and cells. In case of pyrimidine biosynthesis, proteins CTP synthase (PyrG), aspartate transcarbamylase (PyrB) and nucleotide diphosphokinase (Ndk) are quantified to be shared in spores and cells.

### Stress response

*Bacillus subtilis* has to cope with different environmental stresses such as heat, desiccation, fluctuating pH and oxidative stress. All these conditions lead to protein folding stress followed by destabilization of the protein structure^67^. To survive through folding stress and protein aggregation bacteria have developed the chaperon machinery. Molecular chaperones DnaK and DnaJ protect by binding to the unfolded regions of a protein whereas GroEL and GroES provide a suitable environment for proper folding of the protein. Presence of a molecular chaperon machinery along with trigger factor (Tig) and Clp proteins (ClpC, ClpX) in spores may functionally be instrumental for spore resuscitation. For instance, *Streptomyces spp.*, chaperones in the dormant spores help reactivation of the proteosynthetic apparatus and nascent proteins after core hydration^68^. Proteins YqiW and YphP are bacilliredoxins that protect against hypochlorite stress^69^. In addition, transporter proteins YknX, YknZ and membrane protein YknW produced in vegetative cells protect against SdpC (sporulation delaying protein)^70^. In *B. anthracis* petrobactin is also demonstrated to improve sporulation efficiency of cells^71^. Apart from this, the petrobactin binding and importing proteins YclP and YclQ can also help in iron acquisition and/or protect against oxidative stress^72^. In addition to these proteins, the previously mentioned drug efflux pumps may also be important to optimize an environmental stress response.

## Concluding remarks

The one-pot sample processing method along with ^15^N metabolic labelling has for the first time enabled a reproducible, combined cell and spore quantitative proteome analysis of *B. subtilis*. The analysis outlines a relatively modest proteomic adaptation of this bacterium, when as a survival strategy, it completes spore formation. The proteins shared between the cells and spores contribute to the basic set of proteins that are sufficient for life to survive in an otherwise metabolically inactive spore. These proteins include stress survival proteins, proteases, transporters, protein synthesis machinery, metabolic pathway enzymes including those with a key role in the assurance of sufficient energy generation for spore outgrowth and subsequent vegetative cell proliferation.

