## Supplementary figure 1 for "Towards the minimal proteome of life: Proteome profiles of *Bacillus subtilis* vegetative cells and spores"

**Supplementary Figure 1: Relative levels of proteins belonging to the central carbon metabolic pathway in the dormant spores.**

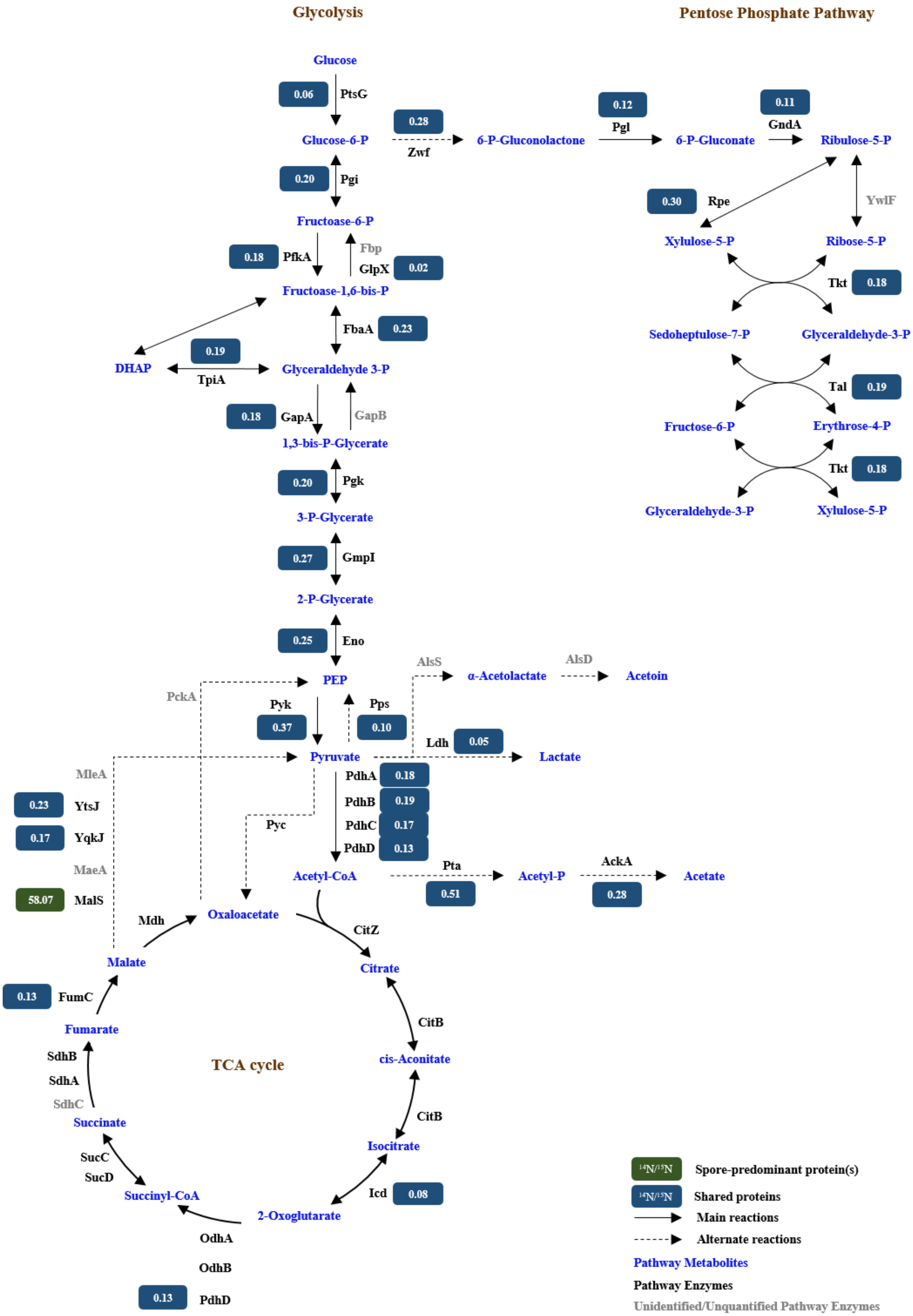

Glycolysis
Pentose Phosphate Pathway
Glucose
0.06
PtsG
Glucose-6-P
0.28
Zwf
6-P-Gluconolactone
0.12
Pgl
6-P-Gluconate
0.11
GndA
Ribulose-5-P
0.20
Pgi
Fructose-6-P
0.30
Rpe
YwlF
Xylulose-5-P
Ribose-5-P
0.18
PfkA
Fbp
GlpX
0.02
Fructose-1,6-bis-P
Tkt
0.18
FbaA
0.23
Sedoheptulose-7-P
Glyceraldehyde-3-P
0.19
DHAP
TpiA
Glyceraldehyde 3-P
Tal
0.19
0.18
GapA
GapB
1,3-bis-P-Glycerate
Fructose-6-P
Erythrose-4-P
0.20
Pgk
Tkt
0.18
3-P-Glycerate
Glyceraldehyde-3-P
Xylulose-5-P
0.27
GmpI
2-P-Glycerate
0.25
Eno
AlsS
α-Acetolactate
AlsD
Acetoin
PckA
PEP
Pyk
0.37
Pps
0.10
Ldh
0.05
Lactate
MleA
Pyruvate
PdhA
0.18
0.23
YtsJ
PdhB
0.19
Pyc
PdhC
0.17
0.17
YqkJ
PdhD
0.13
MaeA
Acetyl-CoA
Pta
Acetyl-P
AckA
Acetate
58.07
MalS
0.51
0.28
Mdh
Oxaloacetate
CitZ
Malate
Citrate
0.13
FumC
CitB
Fumarate
SdhB
TCA cycle
cis-Aconitate
SdhA
SdhC
CitB
Succinate
Isocitrate
SucC
SucD
Icd
0.08
Succinyl-CoA
2-Oxoglutarate
OdhA
OdhB
0.13
PdhD
$^{14}N/^{15}N$ Spore-predominant protein(s)
$^{14}N/^{15}N$ Shared proteins
Main reactions
Alternate reactions
Pathway Metabolites
Pathway Enzymes
Unidentified/Unquantified Pathway Enzymes
